# Using Hi-C and target capture to monitor plasmid transfer in the barley rhizosphere

**DOI:** 10.64898/2026.03.20.713245

**Authors:** Salvador Castañeda-Barba, Thibault Stalder, Eva M. Top

## Abstract

Emergence of multi-drug resistant (MDR) pathogens is facilitated by the mobilization of resistance genes from bacteria in animal and environmental habitats, a process often mediated by plasmids. While fertilization of agricultural soils with manure is hypothesized to serve as a pathway for transferring antimicrobial resistance plasmids to soil and crop bacteria, evidence is limited. In this study, we aimed to determine whether MDR-plasmids from manure transfer in soil, leading to the formation of long-term agricultural resistance reservoirs. To this end, we introduced a known MDR plasmid to agricultural soil where barley was subsequently grown and monitored spread of the plasmid over the course of a growing season (up to 190 days). Our experimental design mimicked conventional agricultural practices at a microcosm scale. A digital droplet PCR approach indicated plasmid transfer in the rhizosphere, which was confirmed by a targeted Hi-C method (termed Hi-C+). This demonstrated transfer of the plasmid to soil bacteria 10 days after barley planting but was not observed afterwards. The new plasmid hosts could not be identified, as plasmid-associated host Hi-C reads were absent from existing databases. This implies these hosts were rare and likely unculturable members of the soil microbiome. Our findings demonstrate that plasmid transfer from manure to soil can occur under conditions reflecting those found in agricultural settings. Furthermore, rare and uncharacterized members of the soil microbiomes may participate in acquiring MDR plasmids from manure bacteria, raising important questions about their role in spreading resistance plasmids.

## Introduction

The spread of antimicrobial resistance among bacteria is a global threat to human health [1]. Mitigating this threat requires understanding of how antibiotic resistance genes (ARG) emerge in pathogens and spread across habitats. Multi-drug resistant (MDR) pathogens are emerging in part due to the horizontal gene transfer mediated by plasmids [2–5], mobile genetic elements that often confer resistance to various antibiotics and can transfer to diverse bacteria [6, 7]. Within habitats, they facilitate rapid bacterial adaptation to changing conditions, such as antibiotic selection [5, 8]. They also facilitate ARG spread between habitats [2].

Farm and feedlot settings are one habitat where the spread of plasmid-mediated antibiotic resistance is particularly concerning. On farms, animals receive antibiotics either preventively or to control disease outbreaks, and in some parts of the world, they are still used as growth promoters [9–11]. In 2013, food animals globally received an estimated 131,109 tons of antimicrobials [12]. This was comparable to the amounts received by humans for the same year [12]. About five years later, in 2017-2018, the Natural Resources Defense Council (NRDC) reported that only 35% of medically important Antibiotics sold in the U.S. were used for human medicine, while the remainder was for cattle, swine and other livestock production [13]. Due to the selective pressure imposed by antimicrobial usage, farm settings are recognized as important sources of antibiotic-resistant bacteria (ARB) and ARG [14, 15]. Furthermore, plasmids that confer resistance to drugs of last resort such as colistin have been found to emerge from this habitat [16, 17]. While farm settings are recognized as important reservoirs of ARG, we still don’t know the trajectories of ARG-carrying plasmids from farms to human pathogens.

ARB and their resistance genes can spread from farms through manure application to agricultural soil [18]. In the US alone, over a billion tons of animal manure are produced each year [19]. Surveys have shown that manure is a reservoir of ARB, ARG and human pathogens [20–24]. Fertilization with manure has been shown to transiently enrich agricultural soils with antimicrobial resistance determinants [25, 26]. Moreover, substantial amounts of ARB and ARG can persist in soil for several months [27–29]. This emphasizes the long-term risk of manure amendment creating ARB and ARG reservoirs in soil.

Creating an ARG reservoir in manure-amended soil requires plasmids to transfer between bacteria and persist in new hosts. We know since the 1970s that a plasmid can transfer from a donor to recipient in soil [30–34]. More recently, the development of new methods has facilitated the monitoring of plasmid transfer to indigenous soil bacteria and identification of those that acquired the plasmid, termed transconjugants [35–37]. These studies demonstrate plasmid transfer from manure to soil bacteria. Additionally, it has been shown that plasmid transfer frequencies can be 10 to 100 times higher in manure-treated versus untreated soil [27, 38, 39]. The same is true for the rhizosphere, the region of soil adjacent to plant roots directly influenced by root secretions [40–45]. Horizontal plasmid spread in manure-amended soil may create ARG reservoirs, which could then enter the food chain through the harvested crops and eventually transfer into human pathogens.

Despite its importance as a pathway for plasmid-borne resistance spread, important knowledge gaps remain. Most previous studies have monitored plasmid transfer in agricultural soil utilizing experimental setups that did not entirely reflect the conditions present in agricultural settings. For example, they used unrealistically high numbers of bacterial plasmid donors (the bacteria carrying a plasmid and transferring it to other bacteria). Additionally, most studies disregarded established agricultural practices that include a time interval between manure application and planting of crops, by introducing the plasmid donor along with the seeds. These approaches make it difficult to extrapolate results to what may be happening in actual agricultural soil and the rhizosphere of crops. Additionally, few studies have been able to identify the soil or rhizosphere bacteria that acquire the introduced resistance plasmids [35, 37, 41]. Therefore, we still don’t know who the key bacterial players are in the dissemination and retention of plasmids from manure.

In this study, we aimed to determine whether MDR-plasmids from manure transfer in soil, leading to the formation of long-term agricultural resistance reservoirs. We monitored the spread of an MDR plasmid in the rhizosphere of barley using a targeted Hi-C approach, Hi-C+, and an experimental setup that followed established agricultural practices. Hi-C is a cultivation-independent proximity ligation method that links adjacent DNA within bacterial cells, enabling the identification of plasmid hosts in microbiomes. We have recently shown that its detection limit in soil can be improved by enriching the focal plasmid DNA through target capture [46]. We found evidence that an introduced *Escherichia coli* plasmid donor can persist in soil for 80 days after inoculation. Avoiding bacterial cultivation and using a native MDR plasmid, we were able to detect plasmid transfer to bacteria in the rhizosphere 10 days after planting of barley. Strikingly, most of the reads belonging to the transconjugants were absent from the databases and metagenomes searched against, suggesting they are uncultured and rare bacteria that may serve as reservoirs and conduits of ARG.

## Materials and Method

### Strains

The donor strain was *Escherichia coli* K-12 MG1655 Nal::gfp [47], carrying IncP-1 plasmid pB10, a self-transmissible broad-host-range MDR plasmid that encodes resistance to tetracycline, amoxicillin, streptomycin, sulfonamides, and mercury ions [48]. The control soil microcosms were inoculated with an isogenic plasmid-free *E. coli* strain. Reference genomes were obtained from a previous study [47, 48].

### Soil, manure, and barley seeds

Soil was obtained from the USDA ARS Northwest Irrigation and Soils research Laboratory in Kimberly, ID (USA). The collected soil was a Portneuf silt loam, which is representative of many cropped fields in Southern Idaho. The soil was sampled from the border between agricultural fields, had a pH of 7.8 and 89 mg/kg of nitrogen (N). The manure was obtained from manure stockpiles at a single dairy in Southern Idaho. The manure had a pH of 9.0 and contained a dry weight of 7.65 and 27.45 g/kg of phosphorus (P) and potassium (K), respectively. Manure and soil were stored at 4 °C until the experiment. Barley seeds, of the spring variety Kardia, were obtained from the Idaho Foundation Seed program.

### Timeline of microcosms, setup, and maintenance

Microcosms were set up in nursery pots with drainage holes: height of 12.9 cm, 15.2 cm diameter at the top, and 10.9 cm diameter at the bottom. In dry weight, each microcosm contained 139.3 g of manure, 630 g of soil and 70 g of greenhouse mix (Pro Mix BX, Quebec, Canada). These microcosms were then inoculated with about 9 x 10^7^ donor bacteria/g, and water holding capacity (WHC) was maintained at 65% throughout the length of the experiment. Additional setup and WHC details are described in Supplementary Methods.

### Microcosm sampling

Bulk and rhizosphere soil were sampled every 10 days starting on day -30, the day of manure and donor strain addition (Fig. 1). From day -30 to 0, only bulk soil was sampled. Barley seeds were planted on day 0, and both rhizosphere and bulk soil were sampled from day 10 until harvest on day 80. Bulk soil sampling continued for an additional 80 days, until day 160. Approximately 3 3 g of rhizosphere or bulk soil were collected at each sampling point (see Supplementary Methods for sampling details).

**Figure 1:**
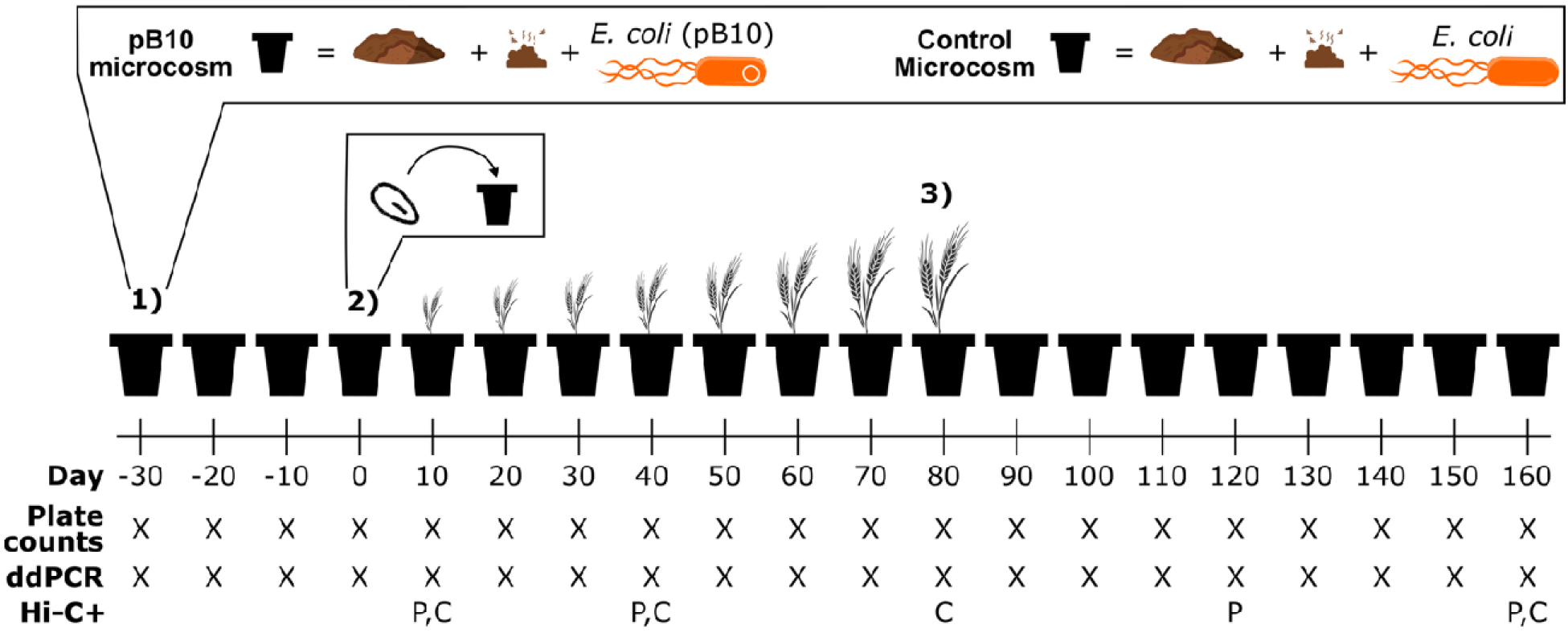
Experimental approach and timeline. The timeline of the experiment is shown below the microcosm containers. Day 0 corresponds to the day when the barley was seeded. Each day depicted is a sampling point. 1) On day -30, we set up microcosms by mixing soil with manure that was inoculated with either the plasmid containing strain *E. coli* (pB10) (called pB10 microcosms), or an isogenic plasmid-free *E. coli* strain (control microcosms). From day -30 to 0, only bulk soil was sampled. 2) On day 0, a germinated barley seed was planted in each soil microcosm. The barley was grown until day 80, when it reached maturity. During this period, both bulk and rhizosphere soil were sampled. 3) The barley was harvested, and sampling of the bulk soil was continued for 80 more days, until day 160. At each sampling point, one control and three pB10-containing microcosms were sampled. The type of data collected at each sampling point is also shown. Due to the high cost, Hi-C+ was only applied soil from one pB10 and one control microcosm on days 10, 40, 80, 120, and 160. Samples to which Hi-C+ was successfully applied are shown at the bottom, with distinctions for successful pB10 (P) and control (C) microcosms.

### Preparation of Hi-C and Hi-C+ libraries

At each time point, 3 g of soil were resuspended in 2% sodium hexametaphosphate (Sigma-Aldrich, St. Louis, MO, USA). Samples were vortexed for 5 minutes and200 μl were reserved for plate counting on selective media. Cross-linked pellets were prepared following Castañeda-Barba et al. [46], using 3 g of soil instead of 5 g and adjusting subsequent volumes accordingly. Cross-linked pellets were stored at -20 °C and ten were shipped to Phase genomics for Hi-C librariey preparation. Eight libraries were successfuly generated and subjected to Hi-C+ target capture enrichment (Fig. 1).

Application of Hi-C+ is detailed in Castañeda-Barba et al [46]. Briefly, Hi-C libraries underwent four PCR rounds before target capture using the myBaits “v5” chemistry and associated v5.01 manual (Daicel Arbor Biosciences, Ann Arbor, MI, USA), using the high-sensitivity protocol. DNA was incubated with plasmid-specific probes and purified to retain probe-bound fragments generating Hi-C+ libraries enriched with pB10-specific DNA.

### DNA extractions

DNA was extracted from soil using the DNeasy Powersoil® Pro Kit (Qiagen, Hilden, Germany). Approximately 0.2 g of soil was used per extraction. DNA was quantified using the Qubit 4 fluorometer with the Qubit™ 1X dsDNA HS Assay Kits (Thermo Fisher Scientific, Waltham, MA, USA) and stored at -20 °C.

### Enumeration of plasmid donor and soil bacteria

During Hi-C preparation, soil suspensions were reserved for plating on selective media to detect the *E. coli* donor. Serial dilutions were plated overnight on lysogeny broth agar supplemented with kanamycin (50 mg L ¹) and nalidixic acid (50 mg L ¹) to detect *E. coli* MG1655 (pB10), or nalidixic acid alone to enumerate plasmid-free *E. coli*. Plates were incubated overnight at 37 °C.

Bacterial densities were also quantified using DNA-based methods. Primers and probes are listed in Table 1. Total bacterial abundance was determined by qPCR targeting the 16S rRNA gene using DNA from each sampling point. Reactions (20 μl) contained Perfecta ToughMix (Quantabio, Beverly, MA, USA), DNA template, and primers and probes at final concentrations of 900 nm and 250 nm. Amplification was performed on a StepOnePlus™ Real-Time PCR system (Thermo Fisher Scientific) using standard cycling conditions. The abundancce of *E. coli* and pB10 was quantified by ddPCR (Table 1). For pB10, we used primers and probe targeting a unique plamid region, described by Bonot and Merlin [49]. *E. coli* primers and probe targeting *gfp* were designed usin Primer3 [50–52]. The ddPCR reactions (22 μl) contained ddPCR Supermix for Probes (Bio-Rad), DNA template, and primers and probes at the same concentrations as above. Droplets were generated on a QX200 droplet generator (Bio-Rad, Hercules, CA, USA), followed by PCR in a C1000 thermal cycler (Bio-Rad, Hercules, CA, USA) using the following cycling conditions: 95 °C for 5 minutes followed by 44 cycles of 94 °C for 15 s and 55.8 °C for 30 s, followed by 98 °C for 5 minutes and finally 4 °C for 15 minutes. The ramp rate at all steps was set to 1 °C/s. The droplets were read using the QX200 droplet reader with default settings.

**Table 1:**
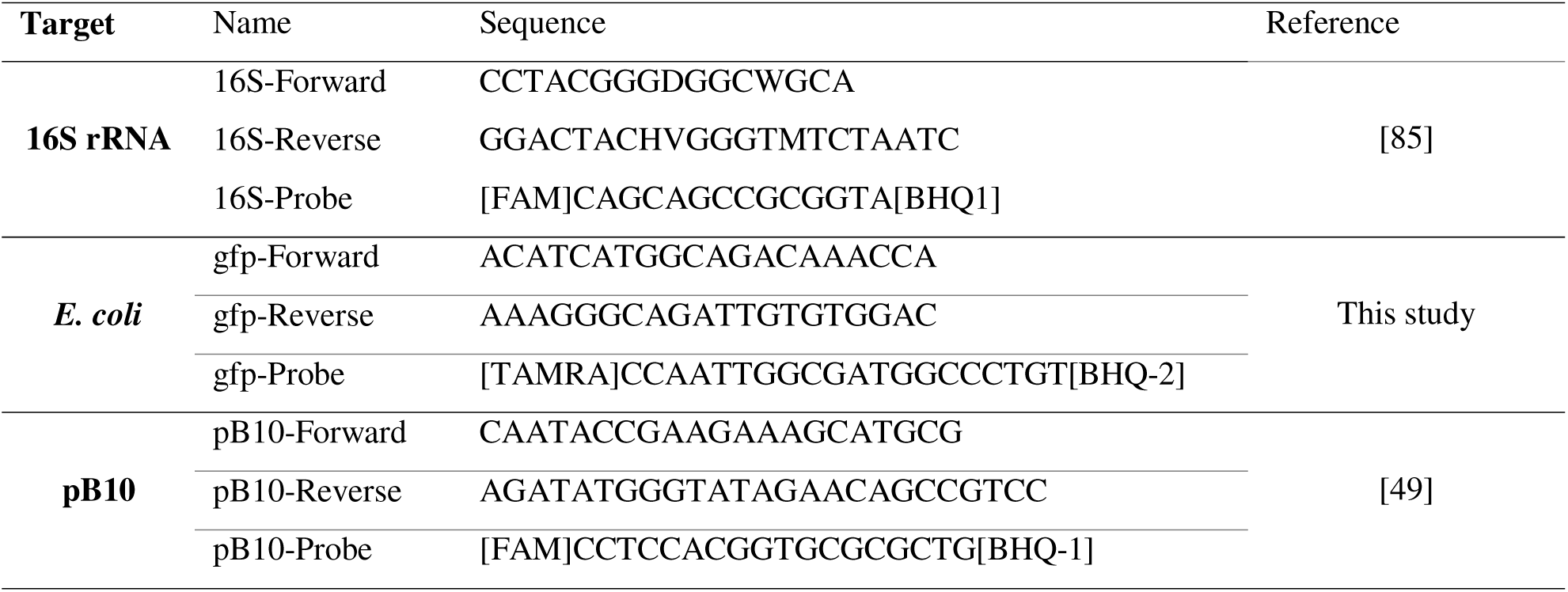
Primers and probes used in this study. Brackets at the start and end of the probe sequences indicate the fluorophore and quencher used, respectively.

### Sequencing of Hi-C+ libraries

Hi-C+ library size selection, quantification, and pooling were carried out by the University of Oregon sequencing core (Eugene, OR, USA). Libraries were sequenced on one lane of NovaSeq 6000 (Illumina, San Diego, CA, USA), using 2 x 150 bp paired-end reads. Read counts per library were as follows; Ctrl_10: 41,038,795, Ctrl_40: 37,530,240, Ctrl_80: 41,577,558, Ctrl_160: 44,354,886, pB10_10: 50,332,918, pB10_40: 35,441,924, pB10_120: 44,774,928, pB10_160: 38,477,065 (Ctrl: control microcosms; pB10: donor-treated mirocosms; _x represent the sampling day).

### Data Analysis

Reads were trimmed and filtered using fastp (version 1.01) with default arguments and minimum length requirement of 50 base pairs [53]. Reads were aligned to reference sequences using BWA-MEM (version 0.7.19) with the -5SPY options [54]. A custom python script (Zenodo DOI: 10.5281/zenodo.18076326) extracted paired reads whereby one pair aligned to pB10 and the other remained unaligned. Unaligned-pB10 reads were retained only if the pB10 segment aligned to the reference genome across its full length with 100% identity. Reads mapping to both the plasmid and donor chromosome were removed to prevent false enrichment signals.

The remaining pB10-associated reads were used to calculate plasmid coverage and unique plasmid bases (Fig. 2). To identify putative pB10 hosts, the unaligned-pB10 reads were extracted and read filtering was carried out. First, reads where the pB10 portion aligned to non-unique regions of pB10 were removed using a python script. Non-unique regions are those on pB10 to which reads from the control also mapped (Fig. 2B). Second, we removed reads that mapped to the *E. coli* and pB10 reference genomes. For removal of a read, two requirements had to be met by its alignment; 1) span over more than 15 base pairs and 2) begin at position one of the read. This second criterion was introduced to address cases where the Hi-C junction, the point at which two fragments of DNA from different sources are ligated, was sequenced. This could result in chimeric reads that partially map to pB10 but are mostly unaligned sequence.

**Figure 2:**
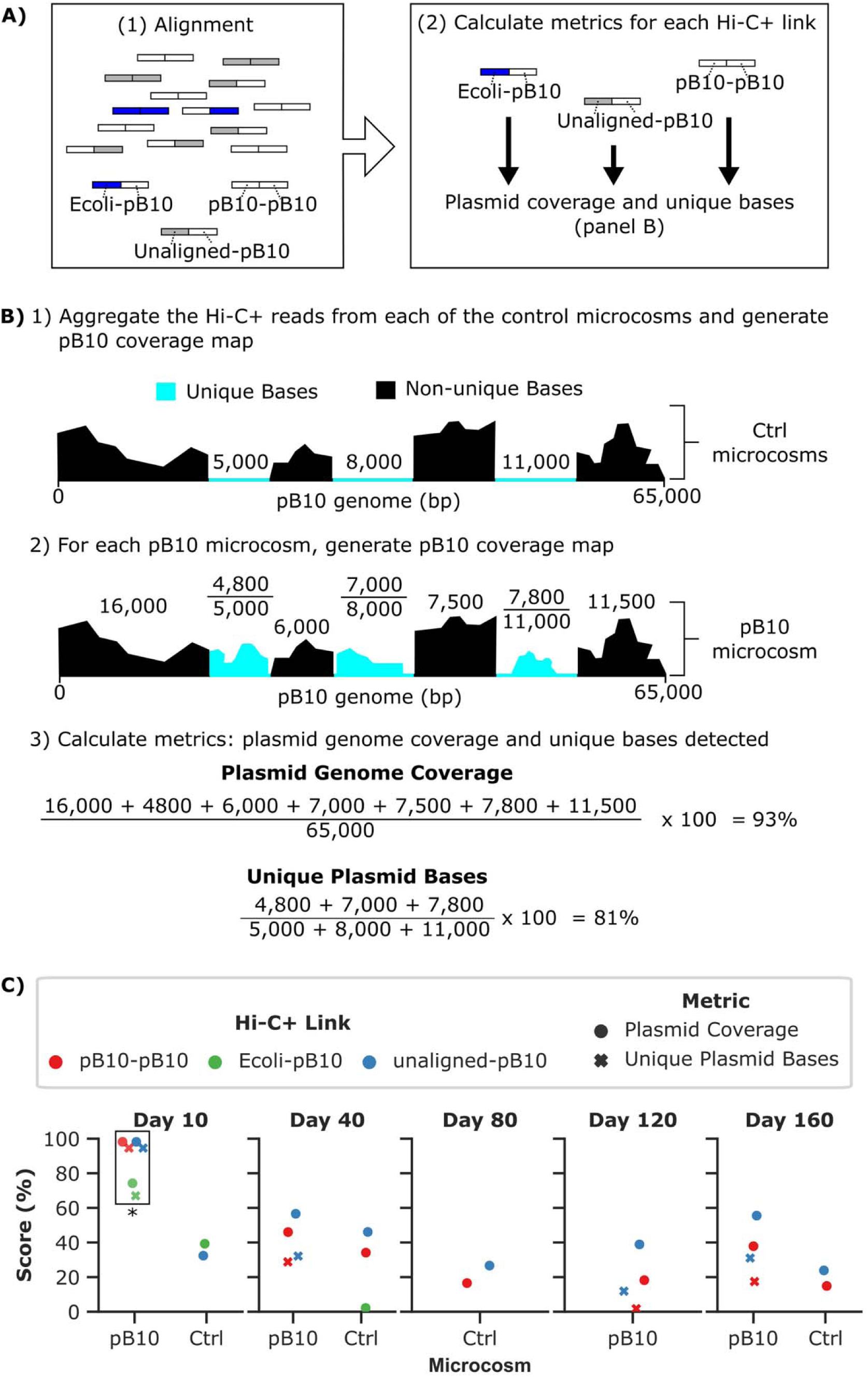
Detection of transfer to soil bacteria, using Hi-C+. A) Overview of data processing pipeline. 1) Reads are first aligned to the *E. coli* and pB10 reference sequences, generating the plasmid-associated Hi-C+ reads depicted. 2) For each of the plasmid-associated reads, two metrics are calculated: plasmid genome coverage and unique plasmid bases. B) Hypothetical calculation of metrics depicting the analysis pipeline. 1) The plasmid-associated reads from each of the control microcosms are first aggregated, and a single pB10 coverage map is generated. Positions in the pB10 genome that are not present in the sequencing data from the control microcosms are denoted as unique bases, shown in blue. The numbers above each region show the length in base pairs. 2) For each pB10 treatment microcosm, a coverage map is generated and is compared against that of the control microcosms. 3) The plasmid genome coverage and unique plasmid bases are then calculated. The calculations shown are based on the pB10 map shown in step (2). Genome coverage is calculated as the percentage of total bases detected in the pB10 microcosm. Unique plasmid bases is calculated as the percentage of unique bases (blue regions) detected in the pB10 microcosm. C) Evaluation of specificity of target capture by comparing metrics calculated for plasmid-associated reads generated from microcosms at each time point. *Metrics observed for the day 10 pB10 microcosm scored close to 100%, suggesting that pB10 DNA segments were successfully linked to putative host chromosomes.

The remaining unaligned-pB10 reads were output to a new FASTA file and used as input for taxonomic classification. First, reads underwent taxonomic classification using Kraken2 (version 2.1.6) with minimum confidence score of 0.1, using the standard database [55]. Taxonkit (version 0.20.0) and csvtk (version 0.34.0) were then used to convert taxids to full taxonomic lineages [56]. Reads unclassified by Kraken2 were then used as input for a BLAST (version 2.16.0) search against the MAG dataset generated from microcosms sampled at days 0 and 10 (see MAG section below). Unclassified reads underwent a search against the NCBI core nucleotide database. BLAST searches were carried out using command line BLAST with default arguments and minimum e-value of 1 x 10^-5^. Lastly, PebbleScout was used to search the remaining unclassified reads against the Metagenomic, WGS, and SRA microbe datasets [57].

### Assembly of metagenome assembled genomes (MAG)

Assembly of MAGs from total soil genomic DNA was carried out for a total of four samples: one control and two pB10-containing microcosms from day 0, and one pB10-containing microcosm from day 10. Total DNA and cross-linked pellets (see above) from each of the four microcosms was sent to Phase Genomics. The total DNA was used for shotgun sequencing while the cross-linked pellets underwent the Hi-C protocol. Phase Genomics then utilized their ProxiMeta^TM^ platform (date pipeline used: September 29, 2023) to assemble metagenome-assembled genomes (MAG) from each sample.

## Results

### Experimental Setup

To study plasmid spread from manure to agricultural soil, we used an experimental setup that mimics realistic processes in agricultural settings. We followed U.S. Department of Agriculture recommendations for both organic and Good Agricultural Practices certification by allowing at least 90 days between manure application and harvest [58]. The details of the setup are shown in Fig. 1. We sought to identify short- and long-term plasmid reservoirs that can be created in soil during an agricultural cycle by monitoring the fate of the focal plasmid in the rhizosphere of barley and in bulk soil after harvest.

### Detection of E. coli and pB10 in bulk and rhizosphere soil

We monitored the presence of the focal plasmid and donor strain at 10-day intervals throughout the 190-day experiment. At each sampling point, abundance of the introduced strain was quantified by plate counting and digital droplet PCR (ddPCR) targeting the *E. coli* and pB10 genomic sequences (Fig. 3). The selective plate count data showed that *E. coli* (pB10) remained in soil for at least 80 days, from day -30 through day 50, after which it was no longer detected in bulk or rhizosphere soil (Fig. 3A). Using ddPCR, *E. coli* DNA was detected until day 110 (Fig. 3B). In contrast, the focal plasmid was detected via ddPCR throughout the entire length of the experiment. No plasmid DNA was detected in the control pots to which plasmid-free donor was introduced, suggesting that the observed signal does not come from indigenous plasmids similar to pB10. These results suggest the plasmid transferred to new hosts, forming a long-term resistance reservoir within agricultural soil.

**Figure 3:**
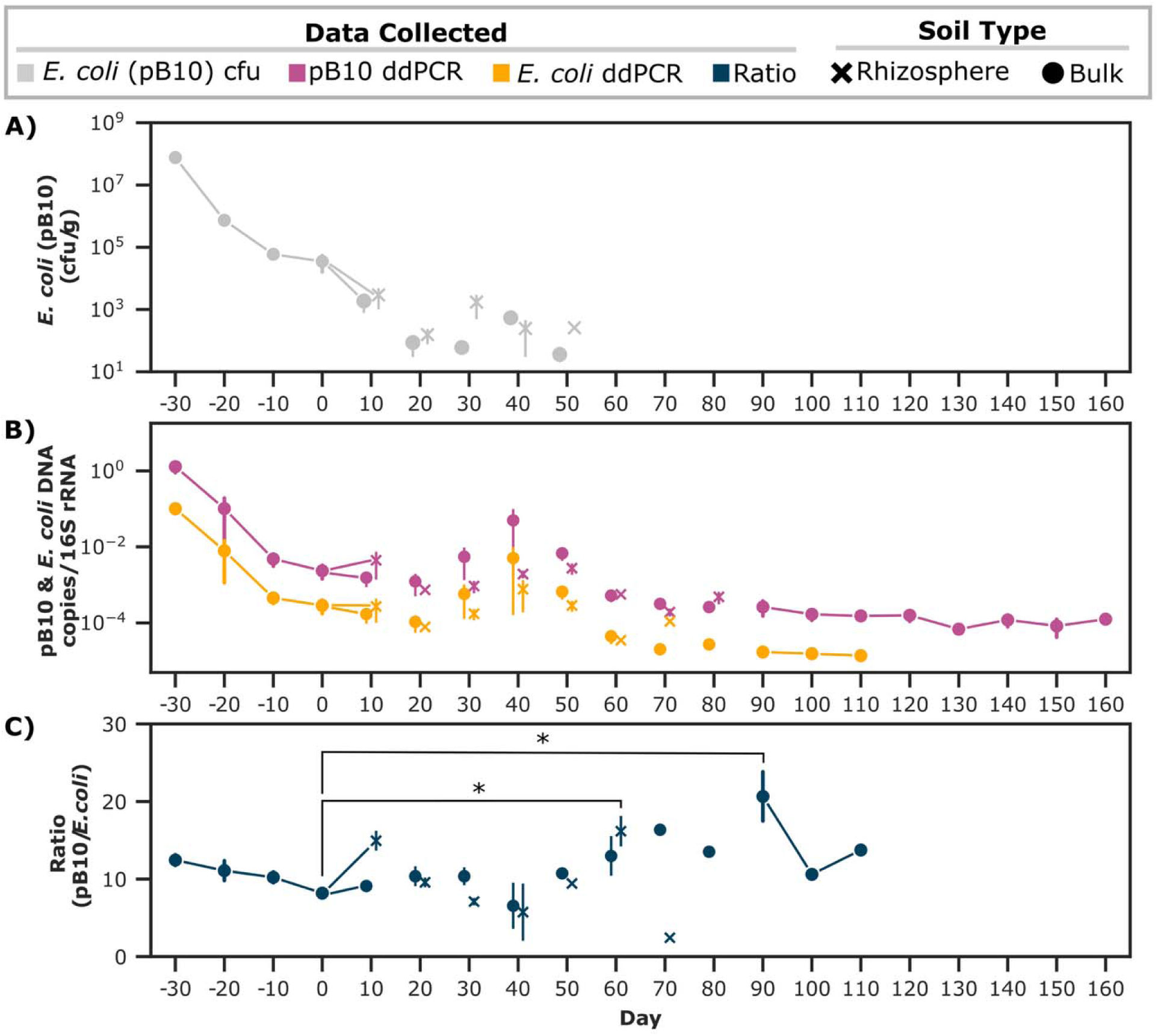
Quantification of *E. coli* and plasmid pB10 throughout the experiment. When the same microcosms were sampled across multiple time points, the data points are connected with a line. In contrast, datapoints obtained by destructive sampling of the rhizosphere are shown a unconnected points. For example, the same three microcosms were sampled from day -30 to 10. Since rhizosphere sampling on day 10 was destructive, a new microcosm was sampled on day 20. A) Detection of the donor, *E. coli* (pB10), via plate counting of soil samples from each tim point on selective media. The donor was no longer detected this way after day 50. B) Relative abundance of *E. coli* and pB10 estimated via ddPCR targeting their respective genomes. Plasmid pB10 was detected throughout the length of the experiment, but *E. coli* was no longer detected after day 110. To account for differences between samples, the detected copies of each target were normalized to the total number of 16S rRNA gene copies present in soil. C) To identify time points where transfer had likely occurred, we calculated the ratio of pB10 to *E. coli*. As pB10 transfers to other soil bacteria, this ratio is expected to increase. Error bars show the standard error of the mean. To identify pairwise differences, we conducted an analysis of variance with post-hoc Tukey’s Honestly Significant Difference tests. Adjusted p-values were used to correct for multiple comparisons, * = p<0.05.

To determine when the transfer of pB10 had likely occurred, we calculated the ratio between the detected copies of pB10 and *E. coli* DNA (Fig. 3C). This approach has been previously used to detect plasmid transfer in environmental habitats [59]. As the number of *E. coli* donors lowers in soil, and the plasmid transfers to other bacteria, the ratio of pB10 to *E. coli* copies is expected to increase. There were two time points at which we observed a significant increase in the ratio of pB10 to *E. coli* DNA, relative to day 0. The first was in the rhizosphere at day 60 and the second in bulk soil at day 90 (Fig. 3C). The increase in ratio suggests that *E. coli* (pB10) likely transferred its plasmid to other bacteria.

### Detection of plasmid transfer to indigenous soil bacteria, using Hi-C+

We next sought to identify new plasmid-host associations using Hi-C+. This method combines Hi-C with a target capture approach to enrich libraries with plasmid-specific DNA [46]. This increases the method’s sensitivity by increasing the proportion of reads linking our focal plasmid to putative hosts in Hi-C libraries [46]. In the paired-end sequence data generated from each Hi-C+ library, we specifically looked for reads that linked the plasmid to itself, to the *E. coli* donor, and to putative new bacterial hosts. These Hi-C+ reads are denoted as pB10-pB10, *E. coli*-pB10, and unaligned-pB10, respectively. Hi-C+ was applied to soil samples from one pB10 and one control microcosm at five time points spread out across the duration of the experiment (Fig. 1). Hi-C+ libraries were obtained for all but two of these ten samples (pB10 treatment day 80, and control day 120). The sampling scheme described here was chosen to spread the data across the length of the experiment and to enable us to identify the short- and long-term bacterial reservoirs of the plasmid.

For each of the Hi-C+ libraries, we first assessed the specificity of the pB10 target capture enrichment by implementing two metrics described by us previously [46] (Fig. 2). The first metric is the percentage of unique bases detected (hypothetical example shown in Fig. 2B). Including control soil microcosms allowed us to assess whether the enrichment was specific to the focal plasmid or was the result of enrichment of similar DNA from soil bacteria. The genes on pB10 with unique portions are listed in Table S1. The percentage of unique plasmid bases is expected to be high when enrichment was specific to pB10, and low when most of the DNA segments were also enriched in the control soil. The second metric is genome coverage. In cases of pB10-specific enrichment, we would expect the pB10 genome to be fully covered. Thus, yielding a high percentage for this metric. Application of these two metrics revealed that enrichment specific to pB10 was only observed with high confidence on day 10 (Fig. 2C, read counts shown in Table 2, column 1). While pB10 was also detected at other time points, the number of unaligned-pB10 reads remaining after stringent filtering to reduce false signals was extremely low, below 400 at every time point aside from day 10 (Table 2, column 3; see next section and data analysis in methods for filtering details). The focal plasmid likely dropped in abundance below that of other plasmids or mobile genetic elements in soil that carry DNA fragments similar to pB10. As a result, the target capture of the Hi-C+ method enriched these similar elements more than, or as much as, pB10, and therefore we could not discriminate pB10 sequences from background noise.

**Table 2:** Count of unaligned-pB10 reads before and after filtering*, pB10 coverage after filtering, and number of filtered reads classified.

| Day | Before filtering | 1<sup>st</sup> filtering | 2<sup>nd</sup> filtering | pB10 Coverage<sup>x</sup> (%) | Classified (N) |
| --- | --- | --- | --- | --- | --- |
| 10 | 119,317 | 6,490 | 6,444 | 25 | 50 |
| 40 | 32,445 | 376 | 375 | 5.2 | 4 |
| 120 | 31,016 | 4 | 4 | 1 | 0 |
| 160 | 37,554 | 87 | 81 | 5 | 8 |

\*The first set of reads that were filtered out overlapped with the control soil. The second set comprised reads with similarity to pB10 or *E. coli*.
<sup>x</sup>Coverage of pB10 conferred by unaligned-pB10 reads was recalculated using only the reads that remained after filtering.

### Identification of putative hosts in soil

With confirmation that plasmid transfer likely occurred on day 10, we next used the unaligned-pB10 reads to identify the putative new hosts of pB10. We first removed unaligned-pB10 reads whose pB10 portion was also present in the control microcosms and could thus be signals from bacteria in soil that carry plasmids or mobile genetic elements with genetic content similar to that of pB10 (Fig. 4, step 3). This reduced the total number of unaligned-pB10 reads from 220,332 to 6,957 (Table 2, 1^st^ filtering). We also removed unaligned reads that partially aligned to the genomes of the added *E. coli* or pB10, as determined by a blast search with cutoffs of 1e-5 evalue and five mismatches. These were removed as they could belong to the inoculated strains but were not labeled as such due to the stringent alignment requirements implemented in previous steps. The remaining 6,904 pB10-unaligned reads (Table 2, 2^nd^ filtering) were then used to identify putative plasmid hosts with higher confidence.

**Figure 4:**
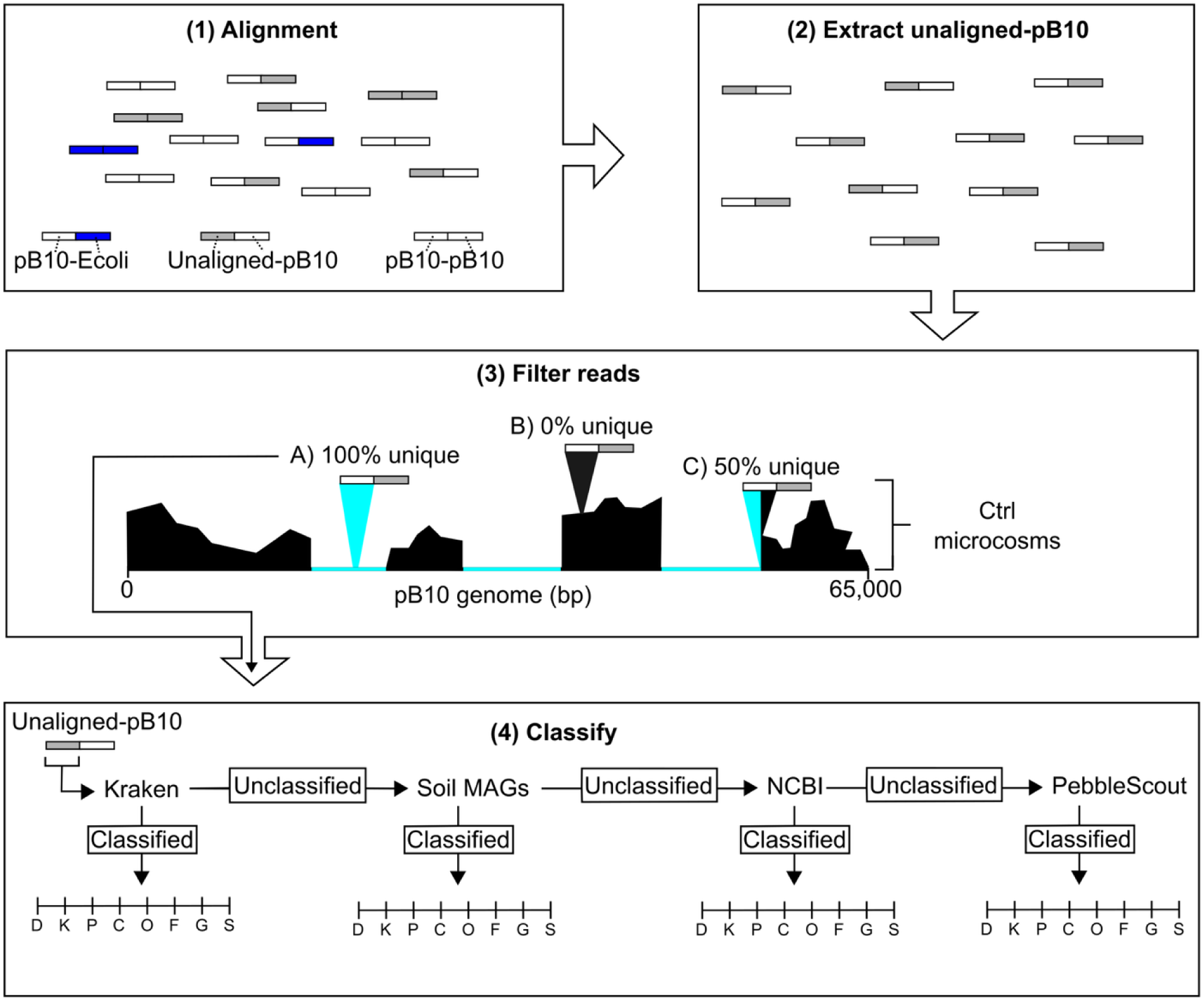
Filtering and classification of reads. 1) Reads were first aligned to the *E. coli* and pB10 reference sequences, generating the plasmid-associated Hi-C+ reads depicted. 2) The unaligned-pB10 reads were extracted from the aligned sequence data. 3) Filtering of reads. First, a pB10 coverage map of reads from the control microcosms was generated. Then, each unaligned-pB10 read from each pB10-treated microcosm was aligned to the control coverage map. The uniqueness of each read was calculated as the percentage of the pB10 portion that aligns to a unique region of pB10, shown in blue. A, B, and C show examples of different uniqueness percentages for unaligned-pB10 reads. Only reads that were 100% unique were retained for classification. 4) The unaligned portion of unaligned-pB10 reads was used to assign taxonomic classification from domain (D) to kingdom (K), phylum (P), class (C), order (O), family (F), genus (G) and species (S). We first carried out a search using Kraken2, a classifier for assigning taxonomic labels to sequencing reads. Reads that could not be classified by Kraken2 were then used to search against a metagenomic dataset generated from soil and rhizosphere samples that were part of this experiment. This dataset consists of 679 Hi-C generated metagenome assembled genomes (MAG) derived from two pB10 and one control microcosm at day 0 and one pB10 microcosm at day 10. This dataset is described in more detail in the methods section. Reads that had no hits to this dataset underwent a BLAST search against the NCBI nucleotide database. A final search was performed for the remaining reads using PebbleScout. After all four searches were implemented, the classification results were aggregated.

To identify the bacterial origin for each unaligned sequence from unaligned-pB10 reads, we sequentially searched against four databases (Fig. 4, step 4). After each search, the reads that were not present in the database searched against were labelled as unclassified and used as input for the next search. Of the 6,904 unaligned-pB10 reads that remained after filtering, 62 were classified using the approach described above (Table 2). The identified genera corresponding to the highest numbers of classified reads (12 and 9, respectively) were *Pseudomonas* and *Escherichia*, but hosts belonging to the alpha- and beta-proteobacteria and a few other classes were also detected (Fig. 5, Table S2). Despite further expanding the search to include MAGs and additional sequencing databases, 98% of the unaligned-pB10 reads remained unclassified. While a small number of reads were detected in PebbleScout datasets, these reads either mapped to raw sequencing data for which taxonomic information was unavailable, or no more than 55% of the read mapped to a provided assembly. This suggests that the potential new host(s) of our plasmid in the rhizosphere were bacterial species whose genomes are not present in existing sequence databases nor in our own soil metagenomes. Thus, while the data strongly suggest that pB10 transferred to bacteria in the rhizosphere, we were unable to identify most of the putative hosts with high confidence.

**Figure 5:**
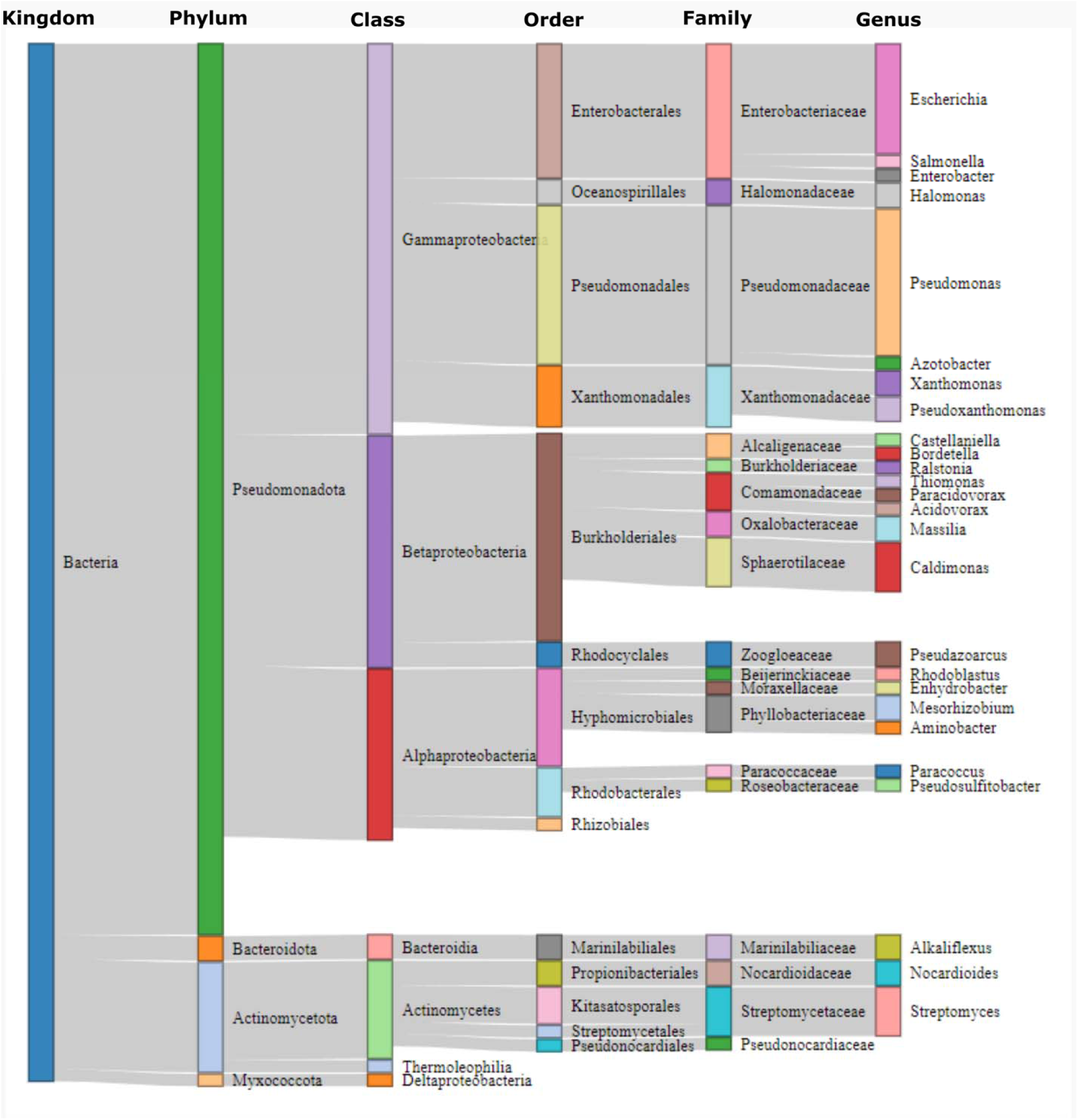
Result from assigning taxonomic classification to the unaligned portion of unaligned-pB10 reads. Only the 62 classified reads are shown here. The taxonomic labels are shown on the top, from Kingdom to Genus.

## Discussion

Plasmids play an important role in mediating the transfer of ARG within and between habitats [16, 17, 60]. To limit the spread of antimicrobial resistance to human pathogens, it is crucial that we identify the bacteria that act as reservoirs for ARG-carrying plasmids [2]. To this end, we monitored the transfer of a MDR plasmid from manure to bacteria in the rhizosphere of barley.

Our experiment closely replicates agricultural practices, which is critical for understanding plasmid spread in natural settings. To our knowledge, no previous study has adhered to these practices as closely as we have. First, we followed agricultural timelines for manure amendment and crop harvesting. Second, our timing of donor inoculation represents a likely scenario for plasmid transfer from bacteria present in manure to rhizosphere bacteria in agricultural settings. A waiting period is typically implemented between the application of manure and the planting of seeds. Thus, a plasmid-host pair present in manure would need to either persist in soil long enough to spread its plasmid in the rhizosphere or transfer its plasmid to soil bacteria before the rhizosphere has developed. Previous studies have not allowed for a time buffer between the inoculation of bacteria and the planting of crop seeds [41]. In some cases, seeds were even soaked in bacterial suspensions, stacking the odds in favor of detecting plasmid transfer [45, 61]. Third, the quantity of plasmid donors added corresponds to a realistic abundance in manure. Depending on the manure and the treatments it undergoes, *E. coli* densities have been reported to range between 10^5^ and 10^7^ cfu/g [62, 63]. While the presence of MDR plasmids in these *E. coli* strains has not been quantified and abundances can vary between manure samples, our inoculation of donors at 10^7^ cfu/g falls within reasonable expectations. Lastly, we monitored the spread of the focal plasmid throughout the entire agricultural cycle, i.e. during barley growth and after harvest. Previous studies that have monitored plasmid transfer in soil or rhizosphere have done so over rather short time periods (hours to days) [37, 41, 43, 64]. While this increases the likelihood of detecting plasmid transfer, it is not informing us about the persistence of the plasmid in agricultural soils over time. Collectively, our approach bridges experimental plasmid transfer studies with agricultural risk assessment.

Throughout the agricultural cycle simulated in our soil microcosms, we observed that the donor remained detectable in soil via plating for 80 days. This is consistent with other studies showing the persistence of *E. coli* in soil over extended periods [65, 66]. Additionally, the higher ratio of plasmid pB10 to donor *E. coli* DNA at days 60 and 90 suggested that pB10 had transferred to other bacteria [59]. On day 60, this higher ratio could be due to replicate-specific dynamics, whereby the focal plasmid was able to transfer and persist in the microcosm sampled at this time point and not in others. On day 90, the detection of transconjugants may be due to mixing of soil carried out on day 80 at harvest time, which may have facilitated the dissemination of nutrients and encounters between plasmid-free and plasmid-containing cells, resulting in detectable transfer. Alternatively, it may have increased the abundance of transconjugants formed earlier in the experiment that were undetectable before day 90. An important note is that ddPCR can also quantify free DNA in the environment [67]. Therefore, other methods are needed to confirm that the plasmid was present in bacteria other than the donor.

To address this important gap, we implemented Hi-C+ to detect plasmid transfer and identify the new host(s). By applying this novel approach to the question of plasmid transfer in soil for the first time, we detected plasmid transfer to new hosts by day 10. However, transfer at later time points was not unequivocally detected, given the background noise from pB10-like sequences in the control soil. The observation of transfer within 10 days of planting germinated barley seeds, and more so in the rhizosphere than in bulk soil, aligns with findings in the literature indicating that the plant rhizosphere environment promotes horizontal gene transfer [42, 43, 64]. A limitation of Hi-C+ is that the target-capture step is a competitive DNA amplification reaction, causing similar DNA fragments in soil to be enriched over the focal plasmid when they are more abundant. For example, plate counts indicated that the donor, *E. coli* (pB10), was still present in the microcosms at day 40 (Fig. 3A), but Ecoli-pB10 Hi-C+ reads were not detected at this time point (Fig. 2C). As a result, not detecting plasmid transfer after day 10 could be the result of non-specific enrichment compromising our detection limit, and does not necessarily suggest that pB10 was no longer present in other hosts.

When using Hi-C+ to identify the bacteria that had acquired pB10 in the rhizosphere, most of the sequences linked to pB10 were absent from existing databases. Only a small subset (∼2%) could be used to taxonomically classify the putative hosts of pB10 (Fig. 5). Regardless, the putative transconjugants detected here have also been identified as transconjugants in previous experiments that introduced an IncP-1 plasmid in soil, i.e., members of the. Alpha-, Beta- and Gamma-*Proteobacteria* [35, 36, 68, 69]. Additionally, several of the genera from the gamma- and beta-*Proteobacteria* identified here overlapped significantly with those detected in the study conducted by Musovic et al [36]. This highlights that these transconjugants could function as general reservoirs of IncP-1 plasmids in soil. Unfortunately, the limited number of classified reads prevented us from confidently determining if these bacteria were truly hosts of pB10.

The large proportion of unclassified reads highlights that more work is needed to characterize soil bacteria. The absence of a significant portion of reads from sequencing databases and from the MAGs of our own soil samples suggests that pB10 was acquired by rare members of the soil microbiome that may be poorly culturable and whose genomes have not yet been successfully assembled. One would expect that the genome sequences of culturable species should be in the databases by now, and that sufficiently abundant species in our soil microbiomes would have been detected in the metagenomes that were prepared from the same soil [70–73]. While low in abundance, rare species are key members of microbial communities that have been shown to perform important ecological functions [74, 75] and receive antimicrobial resistance plasmids [76, 77]. Identifying these rare transconjugants that could be potential ARG reservoirs, will thus require further efforts.

Lastly, we note two key considerations for the implementation of Hi-C+. The first is the probe design. An optimal approach would be to first sequence the habitat of interest and exclude probes that bind to DNA already present. As a result, significant noise would be removed from sequenced Hi-C+ libraries, increasing its sensitivity and specificity. By using probes that completely covered the pB10 genome, we likely amplified similar plasmids of the incompatibility group IncP-1, known to have very similar plasmid backbones and to be present in soil and rhizosphere [78, 79]. The second key consideration is the input DNA for target capture. The Hi-C libraries generated before target capture contained low amounts of DNA, less than 25 nanograms in six of the eight libraries. Due to this low amount of input DNA coupled with the rare target for the probes, output Hi-C+ libraries contained less than 1 ng DNA/μl in every case. Multiple PCR cycles were required for sequencing, likely reducing the percentage of unique reads in each sequenced library. Increasing the input for Hi-C and optimizing DNA yield could improve the success of target capture. With these considerations, we expect that Hi-C+will be a valuable tool for future studies attempting to monitor the fate of plasmids in natural habitats.

In conclusion, in a microcosm setup that mimicked agricultural settings, we found evidence for the transfer of an MDR plasmid from manure to bacteria in the rhizosphere of barley. While we were unable to confidently identify most bacteria that acquired the focal plasmid, our experiment reinforces the notion that ARG transfer via plasmids occurs in this environment. The low abundance of new plasmid hosts, especially beyond 40 days after inoculating the plasmid donor, is encouraging, as it suggests that plasmid reservoirs in soil generated from manure fertilization may be limited. To evaluate whether plasmid transfer from manure to agricultural soil bacteria represents a significant route for AMR spread to human pathogens, it will be pivotal to further improve Hi-C+ along with other methods that enable plasmid host identification (fluorescence-based, epic-PCR, RNA barcoding, etc.) [37, 41, 80–84].

## Supporting information

Description of supplementary materials

## Acknowledgements

This research was supported in part by the intramural research program of the U.S Department of Agriculture, National Institute of Food and Agriculture (NIFA) grant no. 2018-67017-27630 of the United States Department of Agriculture and the Bioinformatics and Computational Biology Program at the University of Idaho in partnership with the Institute for Bioinformatics and Evolutionary Studies (now Institute for Interdisciplinary Data Sciences, IIDS). The findings and conclusions in this publication have not been formally disseminated by the U. S. Department of Agriculture and should not be construed to represent any agency determination or policy. The authors would like to thank the following individuals: Robert Dungan from the USDA ARS Northwest Irrigation and Soils Research Laboratory in Kimberly, ID for help in acquiring soil; Mario De Haro-Marti from the University of Idaho Extension in Gooding, ID for help in acquiring manure; David Hoadley from the University of Idaho Palouse Research Extension and Education Center for help in acquiring barley seeds; Paul Hohenlohe from the University of Idaho for allowing us to use his growth chambers; Tarah Sullivan and Lee James Opdahl from Washington State University for helpful discussions regarding conditions under which to grow barley; Inna Popova from the University of Idaho for lending us soil sieves; and Clint Elg and Erin Mack for assistance in setting up the soil microcosms for this experiment.

## Author Contributions

E.M.T. and T.S. conceived the project; E.M.T, T.S. and S.C. designed the study; S.C. performed the experiments, collected the data, and performed data analysis, and E.M.T and T.S. helped with interpretation and troubleshooting; S.C. wrote the paper. All authors helped revise the paper.

## Data and Materials Availability

All sequencing data pertaining to this project have been made available at the National Center for Biotechnology Information Sequencing Read Archive (PRJNA1395608). Custom scripts used for Hi-C processing, plasmid-associated read extraction, taxonomic classification, and downstream analyses are publicly available via Zenodo (DOI: 10.5281/zenodo.18076326).

