## Supplementary material for "Using Hi-C and target capture to monitor plasmid transfer in the barley rhizosphere": Description of supplementary materials

The supplementary methods is a word document outlining additional methods for soil sampling. Supplementary table 1 is an excel file that outlines the positions on the pB10 genome that are unique (not detected in the control microcosms). Supplementary table 2 is an excel file summarizing the classification of unaligned-pB10 reads.
