## Supplemental methods for "Using Hi-C and target capture to monitor plasmid transfer in the barley rhizosphere"

**Title**

**Determination and maintenance of water holding capacity (WHC)**

Throughout the experiment, the soil was maintained at a predetermined WHC. To know how much water we needed to add to each microcosm, we first identified the weight of the soil at 0% and 100% moisture content. The weight at 0% was identified by placing 25 g of soil mix in a 55 °C oven. It remained there for one week, after which it was weighed again to identify the weight at 0% WHC. Concurrently, 25 g of soil mix was placed on a funnel, and 50 mL of water was added. After 24 hours, the flow through was measured and the amount of water absorbed by the soil was added to its weight. This allowed us to identify the weight of the soil at 100% WHC. Now knowing the weight at 0% and 100% WHC, we were able to determine that our soil mix was initially at 26% WHC.

**Set up of soil microcosms, planting and harvesting of barley**

*1) Set-up of microcosms (Figure 1, step 1)*

*1. A) Mixing of soil with greenhouse mix and addition to microcosms*. Soil for the barley microcosms was first sieved using a 2 mm sieve and subsequently mixed with greenhouse mix at a 90:10 ratio by weight, hereby referred to as soil mix. Due to the clay-like properties of the soil, mixture with greenhouse mix was required for the soil to be conductive to barley growth. Pro-Mix BX Mycorrhizae greenhouse mix was obtained at the University of Idaho and was stored at room temperature until the time of the experiment. After thoroughly mixing, 700 g of soil mix was placed in each microcosm and the mixture was brought up to 50% WHC. We initially brought the soil only up to 50% WHC to allow for additional liquid when adding bacteria and manure to each microcosm. A total of 46 microcosms were set up for the experiment. They were all placed in a Conviron PGC 20 growth chamber that was maintained at 70% humidity and had 16 hr/8 hr light/dark cycles.

*1.B) Addition of bacteria and manure to soil.* The soil mix in each microcosm was maintained at 50% WHC in the growth chamber for seven days before adding the manure and donor strain. This was done to allow time for the soil bacteria to acclimate to new conditions.

*1.B.I) Preparation of bacteria for addition to manure.* Preparation of the bacterial inoculum was initiated three days before adding them to the soil microcosms. Both strains (plasmid-containing and plasmid-free *E. coli*) were first streaked from glycerol stocks (archived at -70 °C) onto Luria Broth (LB) agar plates (LB; Thermo Fisher Scientific, Waltham, MA, USA) with appropriate selection, nalidixic acid (50 mg L^-1^) and tetracycline (10 mg L^-1^) for *E. coli* (pB10) and nalidixic acid alone for the plasmid-free *E. coli*. This selection was used in every culture step henceforth. Subsequently, they were incubated overnight at 37 °C. The next day, a single colony from each plate was inoculated into test tubes with 5 ml LB and grown overnight at 37 °C with shaking at 200 r.p.m. The following day, 30 µL from each overnight culture was aliquoted into 1 L flasks containing 300 mL of LB and the appropriate selection. These cultures were again grown overnight under the same conditions.

The goal of the experiment was to add the *E. coli* strains at a density of about 10% of the total soil bacteria and corresponding to 9x10^7^ donor/g of manure. Due to the large volume of overnight culture that this corresponded to, the overnight suspensions were first concentrated by aliquoting 200 mL at a time into 250 mL tubes and centrifuging them at 5000xG for 20 minutes. The supernatant was removed, and the pellet was resuspended in 100 mL of PBS. This was repeated until all the overnight culture had been concentrated and pooled. From this pooled culture, 42-mL was aliquoted into 50-mL tubes. This volume corresponded to 1.1 x 10^10^ bacteria, which is the number of bacteria to be added to each microcosm.

1.C*) Manure treatment.* The appropriate amount of manure was subsequently set aside for each microcosm, based on the manure treatment used at the USDA research station in Southern Idaho, at a rate of 47.2 tons/hectare or 139.3 g/soil microcosm. This amount was aliquoted in each of 46 50-ml tubes, one for each microcosm.

*1.D) Mixing of manure, bacteria, and soil.* To minimize contamination with plasmid DNA, control microcosms were set up first, followed by the plasmid-containing microcosms. One microcosm at a time, the manure and bacteria were mixed in an autoclaved container. After mixing thoroughly, these were added to a soil microcosm. Along with the manure and bacteria, enough water was added to bring the soil to 65% WHC. The microcosms were subsequently placed back in the growth chamber and maintained for 30 days, at which point the barley seeds were planted.

*2) Planting of barley seeds (Figure 1, step 2)*

To reduce the chances of failed plant growth, barley seeds were pre-germinated prior to addition to soil. Media consisting of 1L of water, 4.1 mL of 0.73 M MgSO_4_, 1 ml of 1.2 M CaCl_2_, and 2 g of phytagel (Sigma Aldrich, St. Louis, MO, USA) was prepared and poured onto petri dishes. The plates were placed at an angle, in a way that the solid media would be deeper at one end of the plate than the other. Three seeds were placed on each plate, with the root node facing towards the deep end. These were placed in the dark for three days. Subsequently, seeds that germinated were used for the soil microcosms. In each microcosm, a germinated seed was placed at a 1-inch depth and was then covered.

*3) Harvesting of barley (Figure 1, step 3)*

Once the barley reached maturity on day 80, it was harvested. The barley plant was cut 1 inch above the soil, leaving the roots behind. To simulate the process of soil tilling, the soil and roots were mixed within each microcosm and placed back into the growth chambers.

**Microcosm Sampling**

*Microcosm sampling - Bulk soil.* For each microcosm to be sampled, a sterile 15 mL tube was inserted into the soil to extract a ‘soil core’. This was done three times, after which the soil was pooled into a 50 ml tube and mixed thoroughly. From this mixture, 0.2 g was used for DNA extraction using the DNeasy Powersoil® Pro Kit. Additionally, 3 g was set aside in a 15 mL tube for Hi-C library preparation, see section on Hi-C library preparation in methods for more details. The remaining soil was stored at -20 °C.

*Microcosm sampling - Rhizosphere soil.* For each microcosm to be sampled, 35 mL of autoclaved rhizosphere buffer (6.33 g L^-1^ NaH_2_PO_4_, 8.5 g L^-1^ Na_2_HPO_4_ anhydrous, pH = 6.5, 200 µl L^-1^ surfactant) was aliquoted into a 50 mL tube. The barley plant was removed from the soil and roots were excised using sterile scissors and placed into the 50 mL tube containing rhizosphere buffer. The tube was then vortexed for 2 minutes, after which the roots were removed using sterile forceps. This process was repeated until roughly 3 grams of rhizosphere soil had been collected. The buffer containing the rhizosphere soil was vortexed to resuspend, and 2 ml were aliquoted into a separate tube to be used for DNA extraction. The remaining buffer containing rhizosphere soil was centrifuged for 10 minutes at 10,000 x g. The resulting pellet was used for Hi-C library preparation. The remaining barley roots were stored at -20 °C.
